## Supplementary data for "A low-complexity linker as a driver of intra- and intermolecular interactions in DNAJB chaperones"

### Supplementary Figures

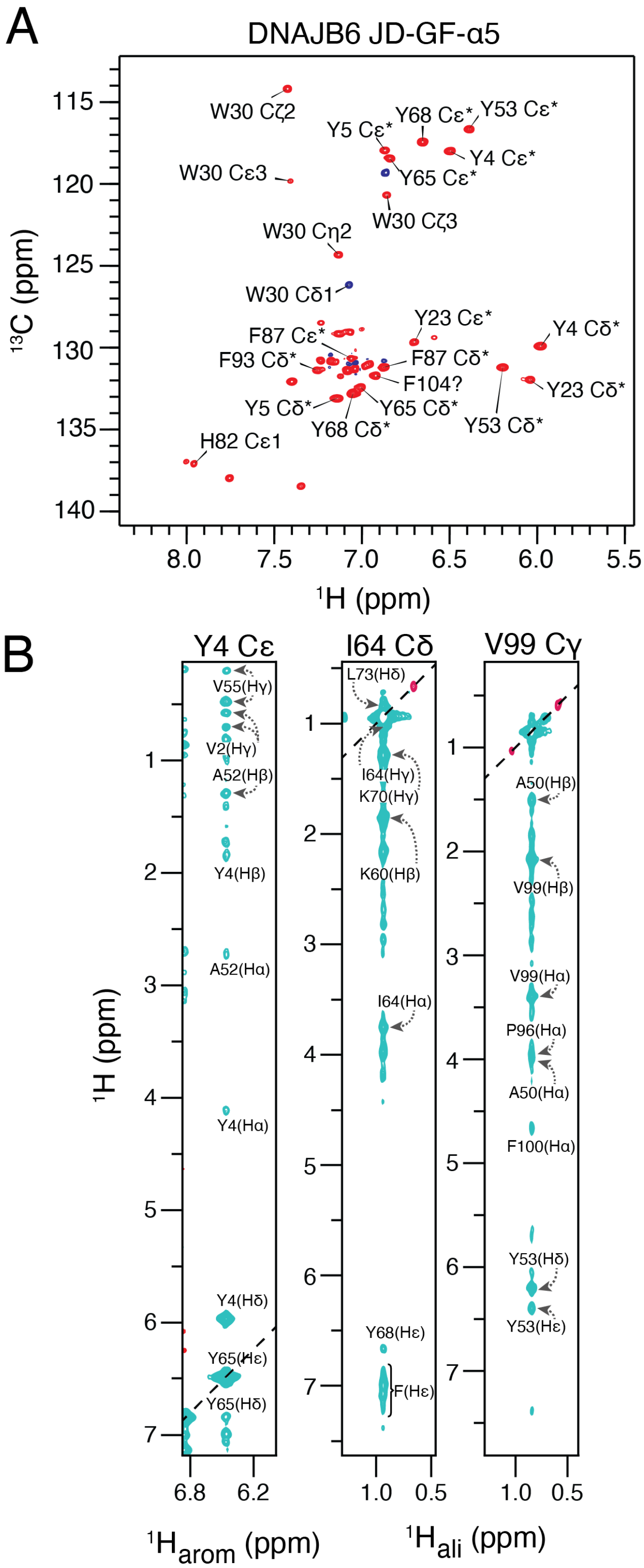


**Supplementary Figure 1: NOE analysis of DNAJB6 JD-GF-α5**. (A) Assigned aromatic $^{1}$H-$^{13}$C TROSY spectrum collected at 800 MHz. (B) Strips from the 3D aromatic (left panel) or 3D aliphatic (middle and right panels) NOESY-HMQC spectra. Both spectra were collected on a sample of 1 mM $^{13}$C, $^{15}$N-labelled DNAJB6 JD-GF-α5.

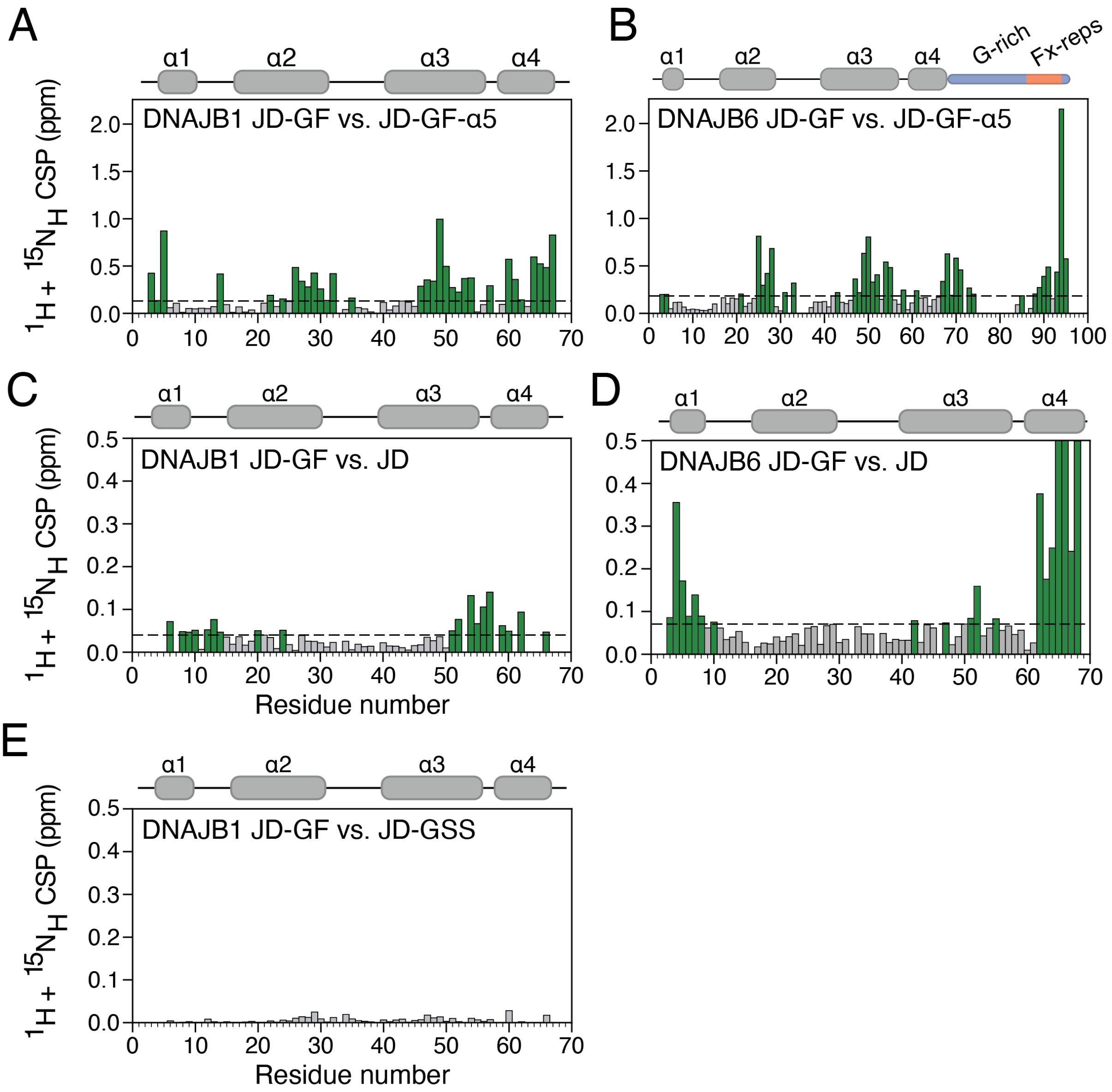


**Supplementary Figure 2: Chemical shift analysis of DNAJB6 and DNAJB1 constructs.** Combined $^{1}H$, $^{15}N$ chemical shift perturbations between DNAJB1 JD-GF and JD-GF-α5 (A) DNAJB6 JD-GF and JD-GF-α5 (B) or DNAJB1 JD-GF and DNAJB1 JD (C) or DNAJB6 JD-GF and DNAJB6-JD (D) or DNAJB1 JD-GF and DNAJB1 JD-GSS (E). Dashed line shows 2 corrected standard deviations, residues that show CSPs above this cutoff (see Methods) are coloured green. (A) and (B) Highlight differences in the NMR spectra resulting from the release of autoinhibition, while (C) and (D) explore the effect of the native DNAJB6/DNAJB1 linker on the chemical environment of JD. (E) Shows that the native GF of DNAJB1 affects the spectrum of JD in the same manner as a completely disorder GSS linker (compare with Fig. 2A).

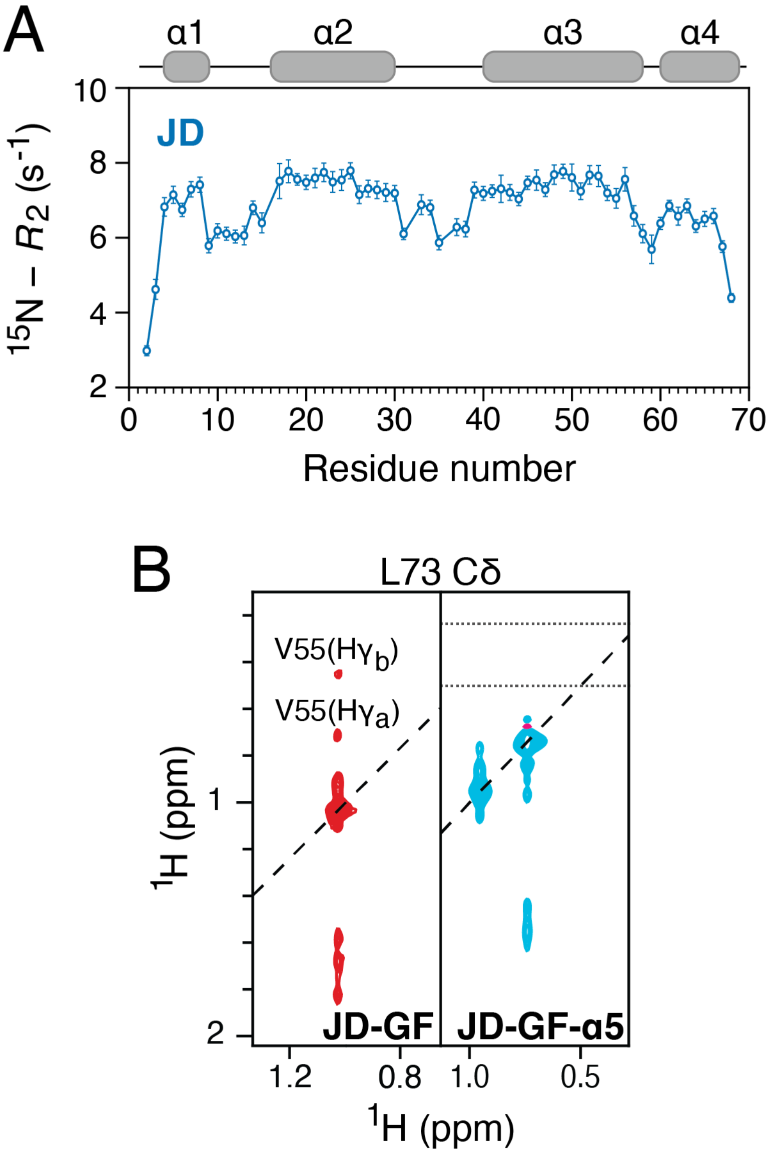


**Supplementary Figure 3: $^{15}$N relaxation rates for JD alone and comparison on NOEs in JD-GF and JD-GF-α5.** (A) $^{15}$N-$R_2$ rates for the isolated JD of DNAJB6 collected on 200 µM $^{15}$N-labelled sample at 600 MHz. (B) Strips from the 3D aliphatic NOESY-HMQC spectra for DNAJB6 JD-GF (left panel) and JD-GF-α5 (right panel). For JD-GF-α5 the chemical shift positions for Val55 Hγ protons are indicated with dashed lines. NOEs were observed between the methyl groups of Leu73 and Val55 for JD-GF but not for JD-GF-α5.

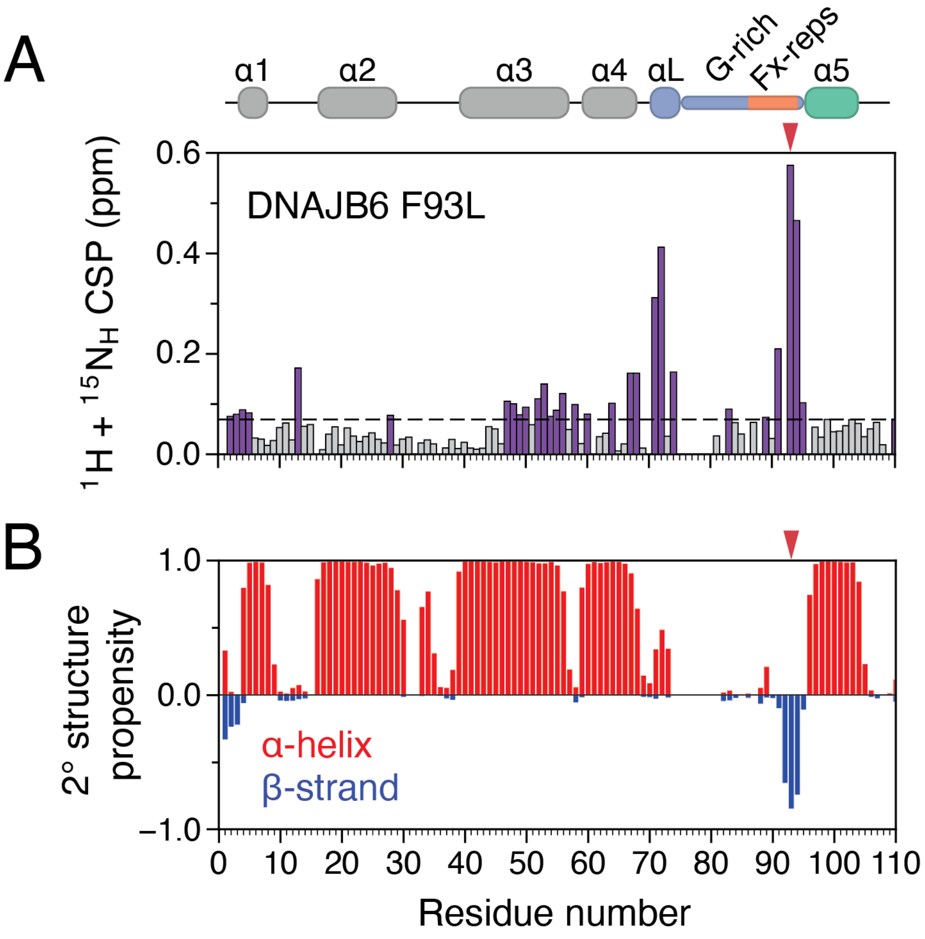


**Supplementary Figure 4: The effect of the F93L mutation on DNAJB6.** (A) Combined $^{1}$H, $^{15}$N chemical shift perturbations between wild-type DNAJB6 and F93L. More widespread CSPs are seen for F93L compared to F91L, with some small changes in the J-domain but not in helix 5. Dashed line shows 2 corrected standard deviations, residues that show CSPs above this cutoff (see Methods) are coloured purple. (B) TALOS-computed secondary structure prediction based on the assigned backbone chemical shifts of DNAJB6 F93L. The position of the mutation is marked with an arrow.

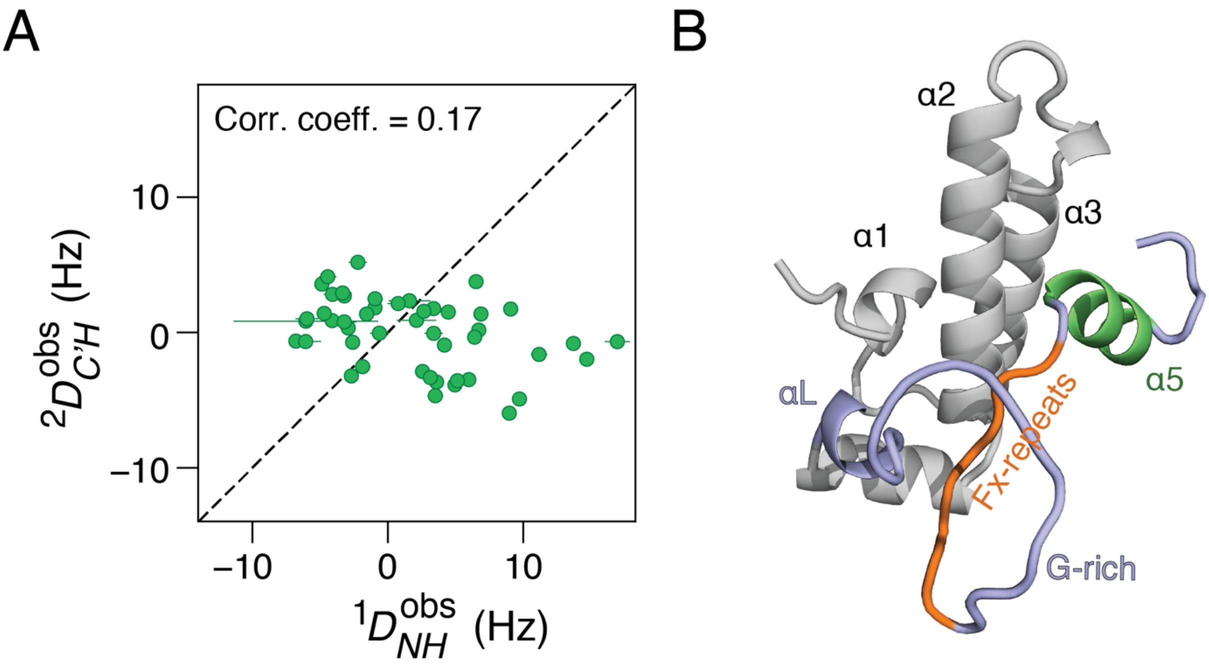


**Supplementary Figure 5: RDC analysis on F91L DNAJB6 JD-GF-α5.** (A) Correlation between the measured $^{1}D_{NH}$ and $^{2}D_{C'H}$ RDC values measured in Pf1 bacteriophage. (B) AlphaFold model of DNAJB6 JD-GF-α5 used to back-calculate the measured RDCs in which residues 70-74 adopt a helical structure (labelled as αL).

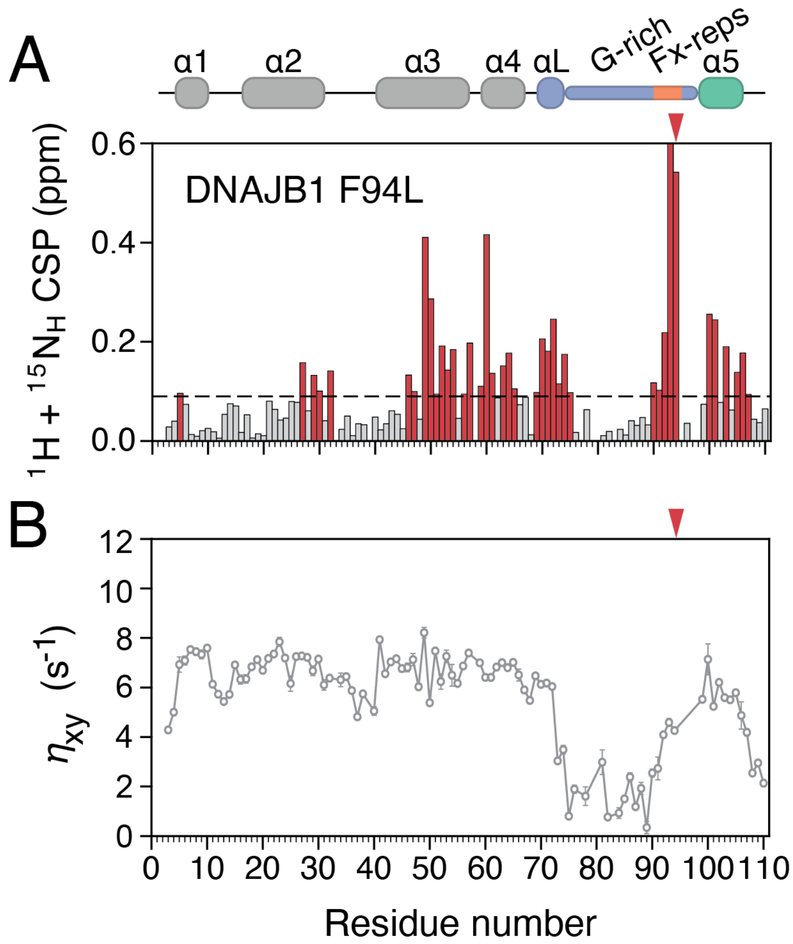


**Supplementary Figure 6: The effect of the F94L mutation on DNAJB1.** (A) Combined $^{1}H$, $^{15}N$ chemical shift perturbations between wild-type DNAJB1 and F94L JD-GF-α5. Dashed line shows 2 corrected standard deviations, residues that show CSPs above this cutoff (see Methods) are coloured in red. (B) Transverse cross-correlated $\eta_{xy}$ rates measured at 600 MHz, on 200 μM DNAJB1 F94L JD-GF-α5 using the pulse sequence of Kroenke et al.[1] The position of the mutation is marked with an arrow.

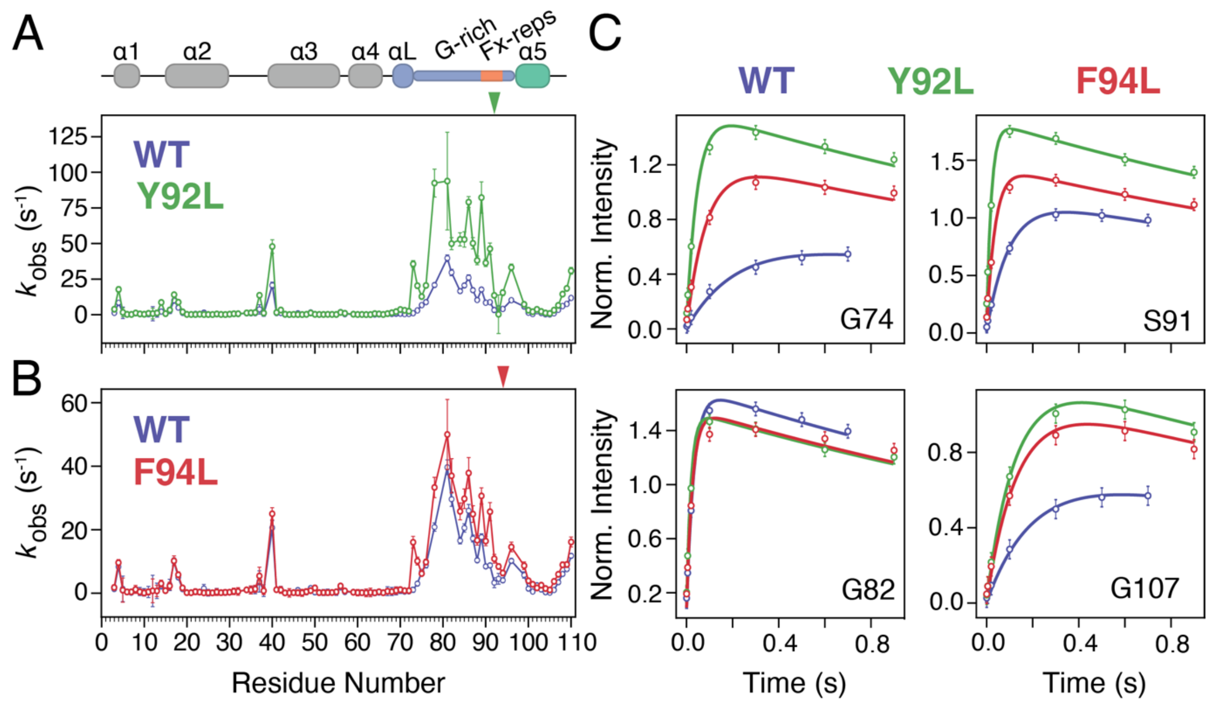


**Supplementary Figure 7: Hydrogen exchange data on DNAJB1 variants.** (A) Comparison of amide hydrogen exchange rates for WT DNAJB1 JD-GF-α5 and Y92L or F94L (B). (C) Selected raw hydrogen exchange profiles measured with WEX-III pulse sequence.[2] The position of the mutation is marked with an arrow.

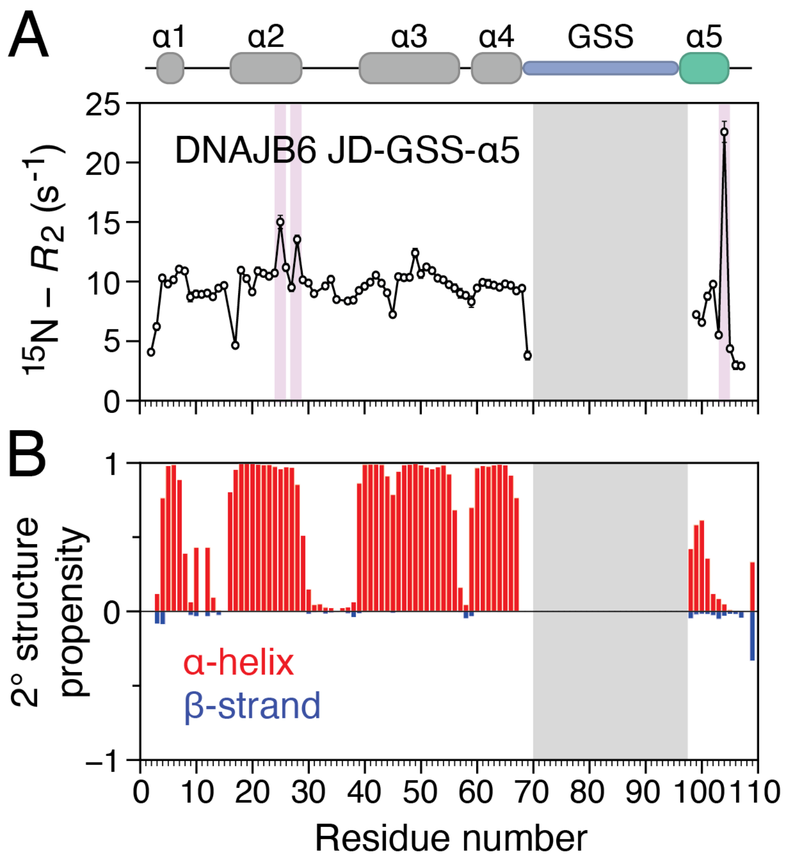


**Supplementary Figure 8: Swapping the GF-linker with disordered GSS destabilises DNAJB6's autoinhibition.** (A) $^{15}$N-$R_2$ relaxation rates of DNAJB6 JD-GSS-α5 construct. Residues with $R_{ex}$ contributions are highlighted in a purple box. (B) TALOS-computed secondary structure prediction based on the assigned backbone chemical shifts of DNAJB6 JD-GSS-α5 (compare with Supplementary Fig. 4B). The GSS resonances could not be assigned as they completely overlap and are noted with a grey box.

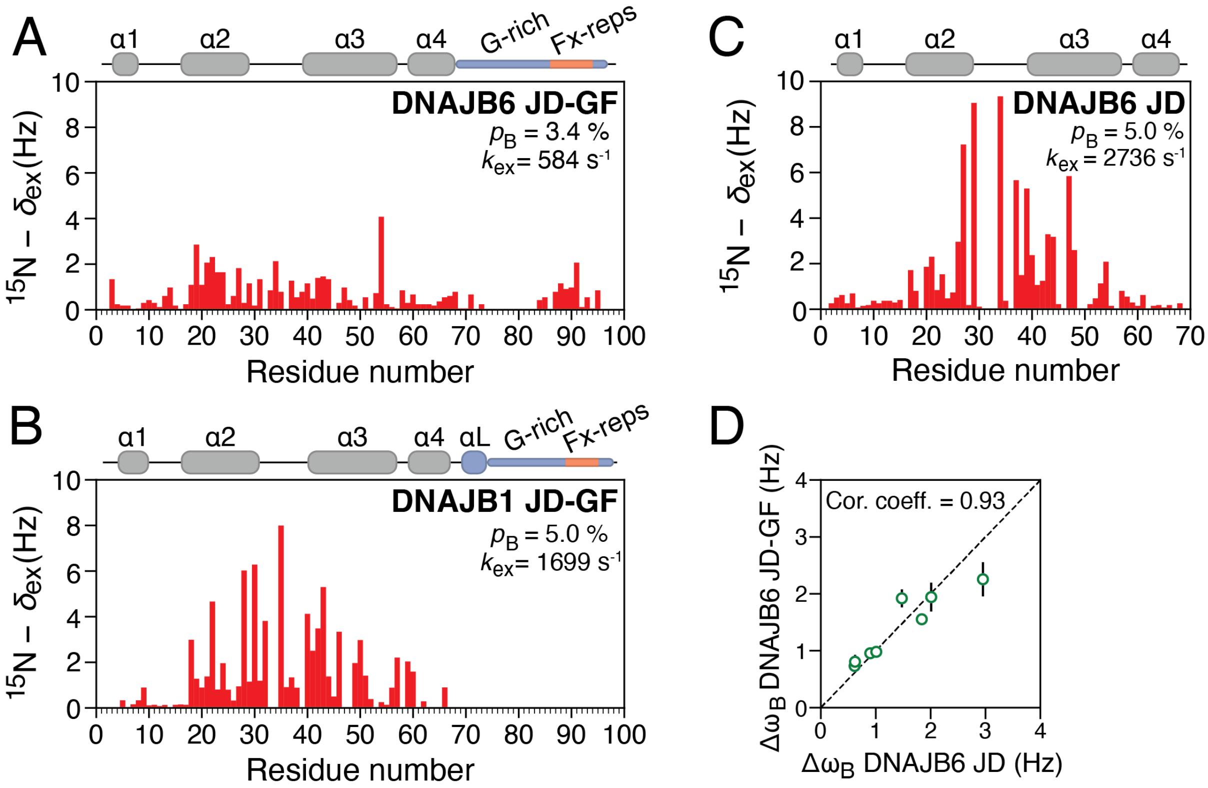


**Supplementary Figure 9: Hsc70 binding to DNAJB6 and DNAJB1.** $^{15}$N-$\delta_{ex}$ values resulting from the binding of DNAJB6 JD-GF (A) or DNAJB6 JD (B) or DNAJB1 JD-GF (C) to full-length Hsc70. (D) Correlation between the fitted chemical shift values for DNAJB6 JD-GF and DNAJB6 JD.

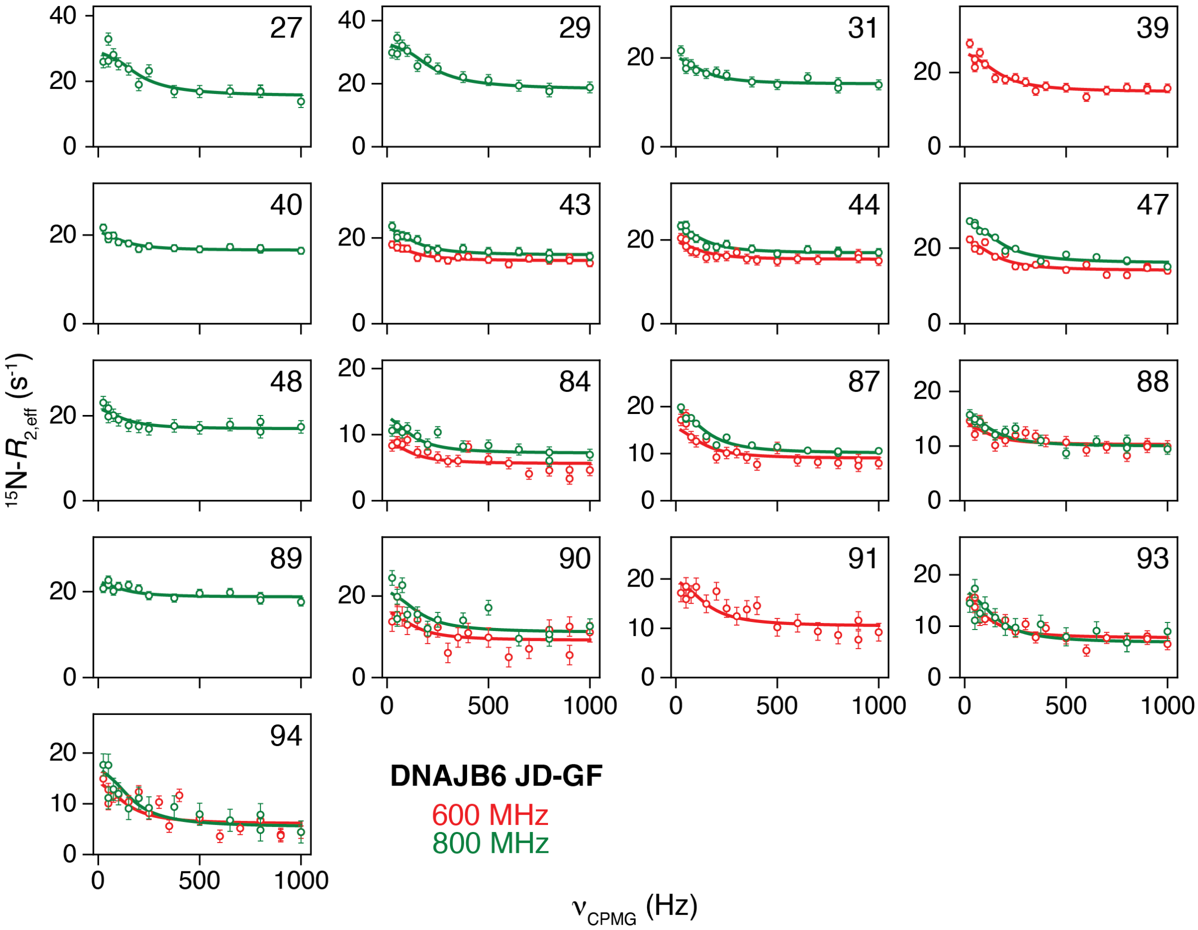


**Supplementary Figure 10: Hsc70 binding to DNAJB6 JD-GF probed by $^{15}$N-CPMG.** In-phase $^{15}$N-CPMG relaxation dispersion profiles of 200 µM NMR-visible DNAJB6 JD-GF in the presence of 20 µM unlabelled Hsc70 recorded at 800 (green) and 600 (red) MHz. The experimental data are displayed as circles, and the continuous lines represent best-fits to the two-state model of Eq. 1 and Fig. 6.

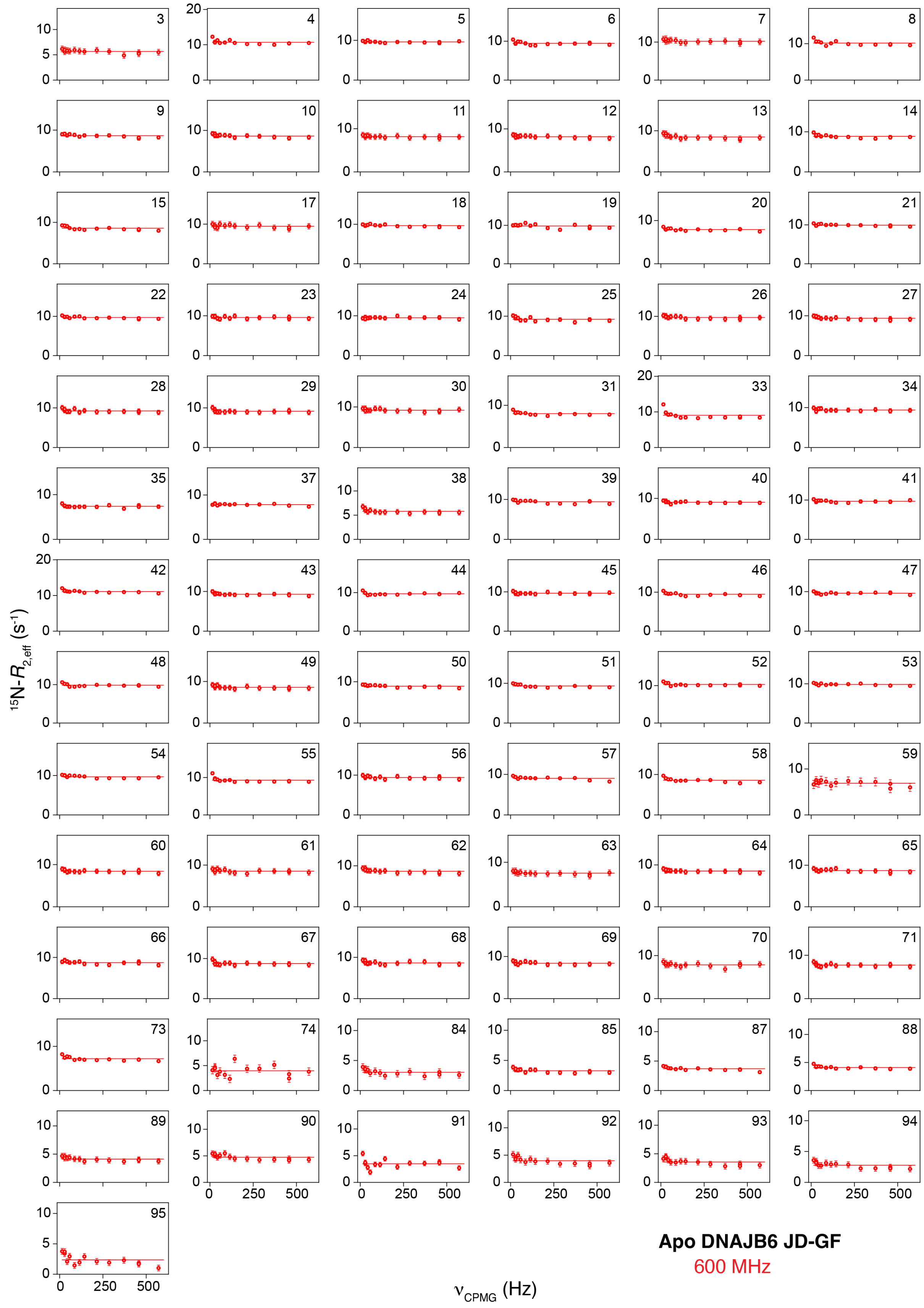


**Supplementary Figure 11: Apo DNAJB6 JD-GF dynamics probed by $^{15}$N-CPMG.** In-phase $^{15}$N-CPMG relaxation dispersion profiles of 300 µM DNAJB6 JD-GF recorded at 600 (red) MHz.

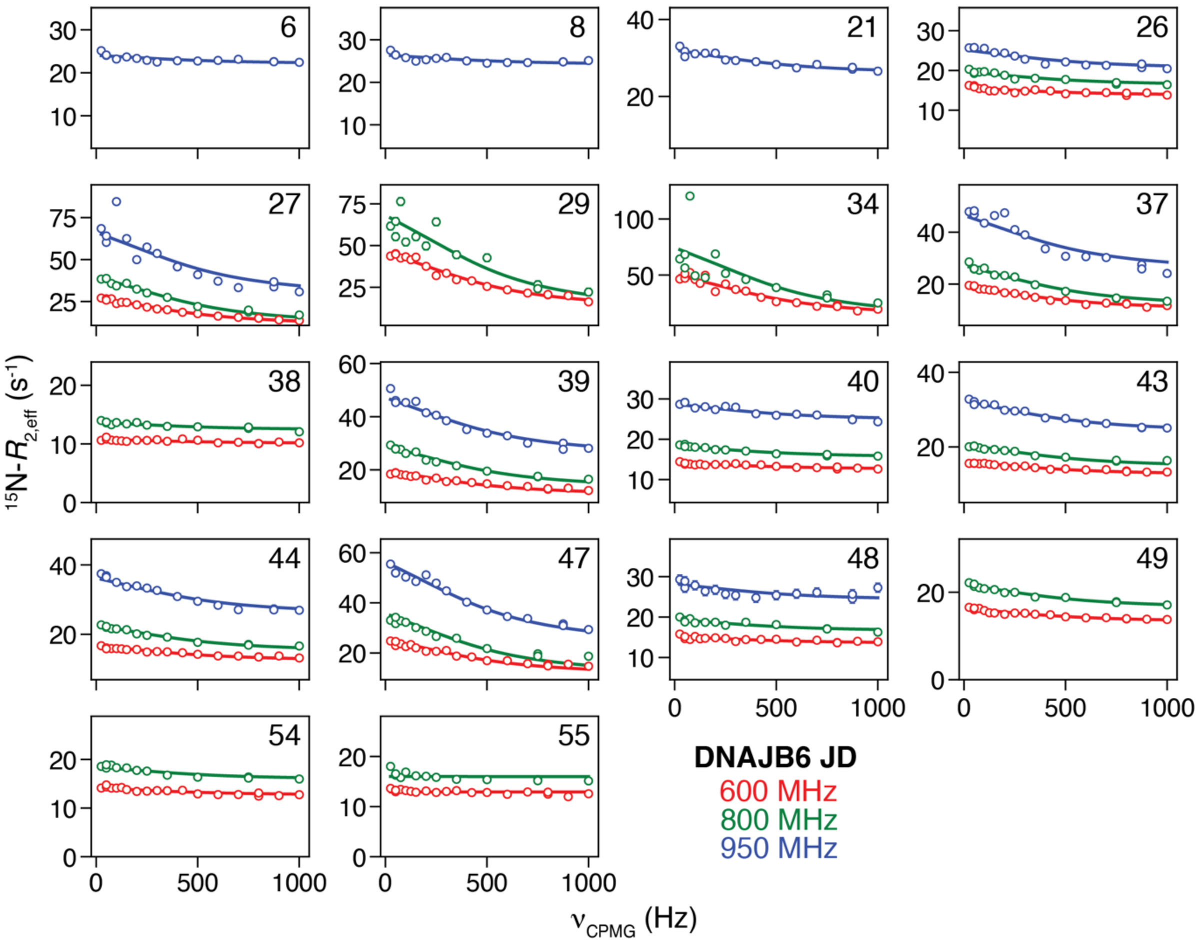


**Supplementary Figure 12: Hsc70 binding to DNAJB6 JD probed by $^{15}$N-CPMG.** In-phase $^{15}$N-CPMG relaxation dispersion profiles of 350 μM NMR-visible DNAJB6 JD in the presence of 35 μM unlabelled Hsc70 recorded at 950 (blue) 800 (green) and 600 (red) MHz. The experimental data are displayed as circles, and the continuous lines represent best-fits to the two-state model of Eq. 1 and Fig. 6.

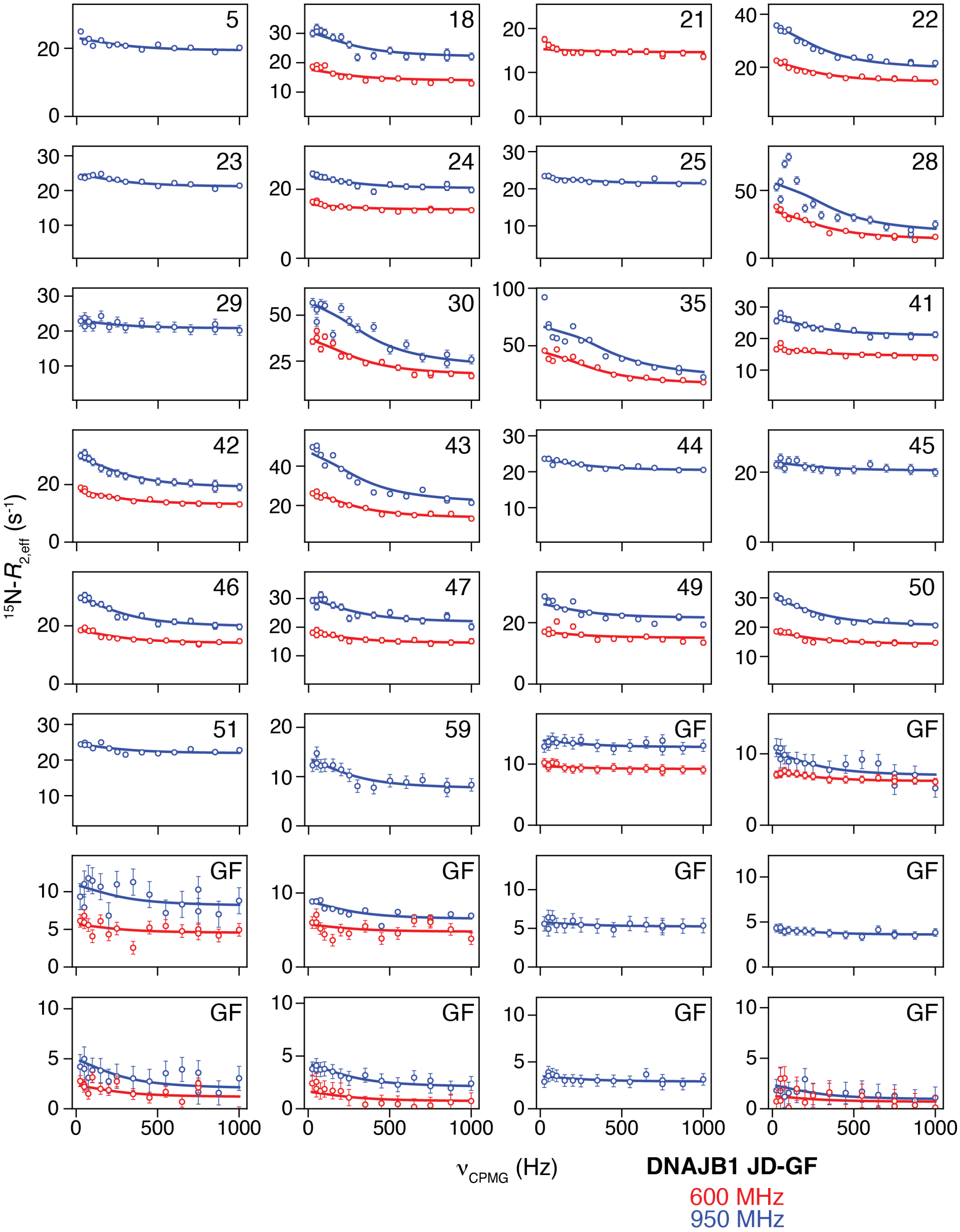


**Supplementary Figure 13: Hsc70 binding to DNAJB1 JD-GF probed by $^{15}$N-CPMG.** In-phase $^{15}$N-CPMG relaxation dispersion profiles of 300 µM NMR-visible DNAJB1 JD-GF in the presence of 30 µM unlabelled Hsc70 recorded at 950 (blue) and 600 (red) MHz. The experimental data are displayed as circles, and the continuous lines represent best-fits to the two-state model of Eq.1 and Fig. 6.

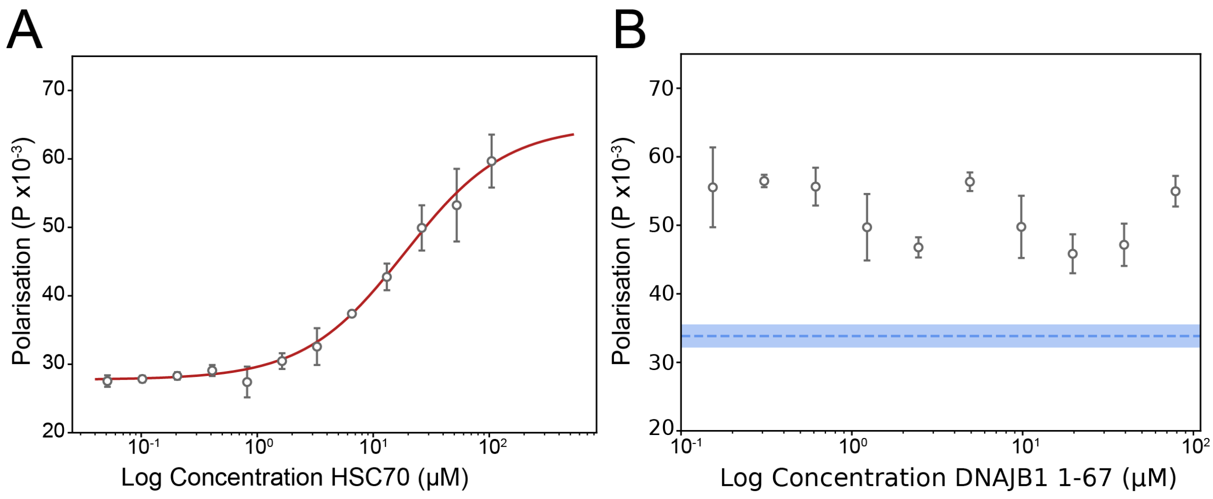


**Supplementary Figure 14: JD binding to Hsc70 measured by fluorescence polarisation.** (A) Titration of Hsc70 (T204A) into fluorescently labelled DNAJB6 JD (300 nM) in the presence of 5 mM ATP. Complex formation was followed by measuring fluorescence polarisation. Error bars show standard deviation for three technical repeats. Red line shows the fit to a four-point logistic curve, which yields a $K_d$ of 18.0 ± 6.5 μM. (B) Competition experiment using DNAJB1 JD. Titration was performed using 300 nM fluorescently labelled DNAJB6 JD in the presence of 40 μM Hsc70 (T204A) and 5 mM ATP. The data show no significant drop in polarisation and hence no displacement of JB6 up to the highest tested DNAJB1 JD concentration, 78.6 μM. Error bars show standard deviation. Dotted blue line shows polarisation measured for free DNAJB6 JD in the absence of Hsc70, blue error band denotes standard deviation for this measurement. All measurements were performed as sets of 3 technical repeats.
